# Investigating the effects of transcutaneous Vagus Nerve Stimulation on motor cortex excitability and inhibition through paired-pulse Transcranial Magnetic Stimulation

**DOI:** 10.1101/2024.07.12.603338

**Authors:** Boscarol Sara, Turchi Letizia, Oldrati Viola, Urgesi Cosimo, Finisguerra Alessandra

**Author notes:** ***Correspondence:** Alessandra Finisguerra. These authors contributed equally to this work and share first authorship (Boscarol Sara, Turchi Letizia).

## Abstract

Transcutaneous Vagus Nerve stimulation (tVNS) has been proposed as a prospective treatment for clinical conditions with altered GABAergic transmission. While possible effects of tVNS on behavioral performance in inhibitory control tasks have been previously reported, neurophysiological evidence showing its effects on GABA-mediated inhibition in the motor cortex is limited. Concurrently, the possible influence of participant’s gender and state conditions remains unexplored. Here, we applied, single- and paired-pulse TMS to the right or the left primary motor in two different groups of participants. We measured corticospinal excitability (CSE), short and long intracortical inhibition (SICI and LICI), cortical silent period (cSP) and intracortical facilitation (ICF) indexes. The measures were taken, in separated sessions of a within-subject design, at baseline prior to tVNS and after delivering active and sham tVNS in the Cymba conchae of the left ear. To exploit state dependent effects and assess the role of tVNS in motor learning, tVNS was applied, during the execution of a computerized visuomotor task. In the left TMS group, we observed better visuomotor performance during active than sham tVNS, regardless of participant’s gender. Interestingly, in both groups, we found a specific increase of SICI, which is mediated by GABAa activity, after active compared to sham-tVNS and baseline evaluations, which was specifically limited to female participants. No effects on CSE, ICF or GABAb-mediated intracortical inhibition indexes were observed. The results show specific effects of tVNS on motor learning and GABAa-mediated motor inhibition, providing supportive evidence for the application of tVNS as an alternative and coadjuvant treatment for disorders featured by altered inhibition mechanisms.

## INTRODUCTION

Transcutaneous vagus nerve stimulation (tVNS) is a relatively new non-invasive alternative to its invasive counterpart (VNS) to increase vagus nerve activation. It consists in the application of weak electrical current to the sensory afferent fibres of the auricular branch of the vagus nerve (ABVN) in the outer ear. From there, the current targets the nuclei of the nerve located in the brainstem, particularly the Nucleus of the Solitary Tract and the Locus Coeruleus (Frangos et al., 2015; Yakunina et al., 2017). Via these projections, tVNS modulates the activity of cortical and subcortical structures (Badran et al., 2018; Dietrich et al., 2008; Kraus et al., 2007; Rajiah et al., 2022; Yakunina et al., 2017).

Overall, tVNS is hypothesized to mimic VNS effects (Assenza et al., 2017) and to modulate various neurotransmitters systems, mainly but not exclusively including the Norepinephrine-(NE) and the γ-aminobutyric acid-(GABA) systems (Farmer et al., 2020). These neurotransmitters play crucial roles in synaptic transmission, neuronal excitability, and neuroplasticity (Brunoni et al., 2008), suggesting that tVNS may modulate specific neural pathways (Badran et al., 2017, 2023; Baig et al., 2022; Groves & Brown, 2005; Hulsey et al., 2019). Considering its potential, several studies have suggested that tVNS could be used to treat diseases characterized by neurotransmitter imbalances, such as epilepsy (Bauer et al., 2016), depression, anxiety and neurodevelopmental disorders (Albin & Mink, 2006; Dale, 2017; Draper et al., 2014; Del Campo et al., 2011; Edden et al., 2012; Jin & Kong, 2016), as well as for rehabilitating motor deficits in stroke patients (Badran et al., 2023; Baig et al., 2022; Capone et al., 2017).

Despite the growing interest in its therapeutic potential, the mechanisms of action of tVNS are not yet fully understood. Since increased GABA levels are linked to improved response inhibition (Boy et al., 2010; Quetscher et al., 2015), behavioural studies demonstrating an enhancement in participants’ performance in inhibitory control tasks during tVNS provided indirect evidence of the possibility to increase GABA activity via tVNS (Beste et al., 2016; Jongkees et al., 2018). However, the direct psychophysiological effects of tVNS on cortical GABAergic inhibition are not yet clearly understood (Herr et al., 2024).

Cortical GABAergic inhibition can be investigated non-invasively by using paired-pulse (pp) and single-pulse (sp) Transcranial Magnetic Stimulation (TMS) protocols. More specifically, pp-TMS protocols consist in the application through the same coil of two TMS pulses at different inter stimulus interval (ISI) and intensities. Pp stimuli applied at very short ISI (1-5 ms) allow measuring short-interval intracortical inhibition (SICI), resulting from the reduction of the response to a test supra-threshold stimuli by a sub-threshold conditioning stimulus (Di Lazzaro et al., 1998; Ilić et al., 2002). At longer ISI (50-200 ms), it is possible to assess long interval intra-cortical inhibition (LICI), consisting in the reduction of the response to a test supra-threshold stimuli by a supra-threshold conditioning one (Valls-Solé et al., 1992). Importantly, evidence from pharmacological studies suggested that SICI and LICI are mediated, respectively, by GABAa-ergic and GABAb-ergic inhibition (Di Lazzaro et al., 2000; McDonnell et al., 2006; Werhahn et al., 1999). Notably, pp-TMS protocols allow also assessing Intracortical Facilitation (ICF), in which the response to a supra-threshold stimulus can be facilitated when paired with a preceding subthreshold stimulus at an intermediate ISI (e.g., 10 ms), and this process is supposedly mediated by noradrenaline (Herwig et al., 2002). Lastly, inhibitory GABAb-ergic circuits can be probed through sp-TMS protocol applied during muscle contraction by measuring the duration of the cortical Silent Period (cSP), which consists in the suppression of electromyographic (EMG) activity after a supra-threshold sp stimulus (Tergau et al., 1999; Werhahn et al., 1999).

These indexes have been exploited to probe the effect of tVNS on cortical inhibition. In particular, following the findings of an increase in SICI after VNS in patients with epilepsy (Di Lazzaro et al., 2004), studies have searched for a similar SICI increase triggered by non-invasive tVNS. While a seminal study confirmed this effect in a small group of healthy participants (Capone et al., 2015), and it has been widely taken as a proof of the similarity of tVNS and VNS effects (Farmer et al., 2020), more recent studies have provided contrasting results (Midden et al., 2023; Mertens et al., 2022; Gerges et al., 2024; Yun et al., 2024). These studies, however, varied in terms of the site and protocol of stimulation, which may trigger more or less optimal vagal activation, the control stimulation, participants’ gender and the laterality of the recorded corticospinal excitability. Furthermore, whether tVNS affects other indexes of cortical inhibition has been underexplored. Finally, while it has been largely demonstrated with other non-invasive brain stimulation techniques that the brain responses to stimulation are state dependent (Silvanto et al., 2007), this has not been investigated for tVNS (Farmer et al., 2020).

To address these issues, here we aimed at assessing the effect of tVNS on GABAa and GABAb mediated cortical inhibition on a sample of male and female participants. In particular SICI, LICI and cSP were assessed as measure of cortical inhibition. Furthermore, corticospinal excitability (CSE) and ICF were measured, to exclude unspecific effect on cortical activation (see Yun et al., 2024). Importantly, tVNS was delivered when participants were engaged in a visuomotor task to control for possible state dependent effects of tVNS on cortical inhibition. This also allowed us to test an effect of tVNS on visuomotor abilities. Lastly, to explore whether the effects of tVNS were actually limited to the contralateral (right) motor cortex or whether they generalized also to the left hemisphere, TMS was applied both to the right and to the left motor cortex, in a between-subject design. In keeping with the findings of Capone et al. (2015), we expected left-ear tVNS to increase intracortical inhibition, at least that mediate by GABAa receptors (i.e. SICI), but not CSE and ICF, and that this effect should be associated with improved visuomotor performance. However, considering the variability of previous findings, we expected that these effects should be moderated by the laterality of the probed motor cortex and by the gender of the participants, with greater effects in female participants and for the contralateral right motor cortex.

## Materials and Methods

### Participants

A total of 49 healthy participants (age range 22.469 ±3,803 years, 29 females) took part in this study. We determined the required sample size for each group in our mixed design (numerator df = 2) by using the G*power software (Faul et al., 2009) with the “as in SPPS option”, setting alpha level at 0.05 and the desired power at 80%. A large effect size of f (U) = 0.52 was estimated based on the resulting effect sizes of previous studies investigating modifications of SICI in male and female participants after left tVNS (Capone et al., 2015; Gerges et al., 2024; Mertens et al., 2022; Midden et al., 2023).

After providing an overview of the study, including general information about TMS and tVNS, all participants, who remained naïve to the experimental hypothesis and to the conditions of stimulation throughout the whole experimental sessions, gave written informed consent for participation. The study was approved by the local ethics committees (Comitato Etico I.R.C.C.S. Eugenio Medea. Prot. N. 66/22) and conducted in accordance with the Declaration of Helsinki. Prior to the experiment, the participants were screened to evaluate their suitability for the application of TMS and tVNS (Farmer et al., 2020; Rossi et al., 2009). None of the volunteers had a history of neurological disorders, brain trauma, or a family history of epilepsy or cardiac problems. Furthermore, all participants were free from medication intake, presented normal or corrected-to-normal vision and were right-handed, as assessed by the Standard Handedness Inventory (Briggs & Nebes, 1975). Only after completing the whole experiment, participants received information about the experimental hypothesis.

### General procedure

In this study we used a randomized, single-blind sham-controlled mixed design. Participants were randomly assigned to the left or to the right TMS group, receiving, respectively single pulse and paired pulse TMS over the left or the right primary motor cortex. For each group, the experiment consisted of three sessions, which lasted approximately 2 hours each. The sessions were structured as follows (see Fig. 1). In the first session (baseline day) a TMS protocol (detailed in the **TMS protocol** section) was applied to assess the individual baseline values of corticospinal excitability, cortical inhibition and facilitation at rest. For both groups, in the following two sessions (active tVNS and sham tVNS day, see the **tVNS** session), participants received 1 hour of either active or sham tVNS during the execution of a Visuomotor task. To ensure sufficient activation of the vagus nerve during the task, participants started the visuomotor task 30 minutes after the onset of the stimulation. Thus, in the first 30 minutes of each session, to control for their activation in a rest condition and to ensure comparability across participants and sessions, they were invited to read a book. Then, they performed the visuomotor task lasting about 30 minutes (see the **Visuomotor task** section). Soon after the end of tVNS, the same TMS protocol they underwent in the baseline day was applied. The safety of tVNS was checked throughout the delivery of tVNS, while the tolerability and the sensations induced by either tVNS or TMS were assessed at the end of each stimulation day (see the **Safety and tolerability evaluation** section). The sessions were scheduled at least 24h apart to ensure a washout period and with a maximum interval of 3 days from one session to the other (from session 1 to session 2: mean= 2.27 days, SEM =0.22 days; session 2 to session 3: mean =1.94 days, SEM= 0.25). The baseline session was always administered as the first session, while the order of the active and sham sessions was randomized between subjects and balanced across groups.

**Figure 1:**
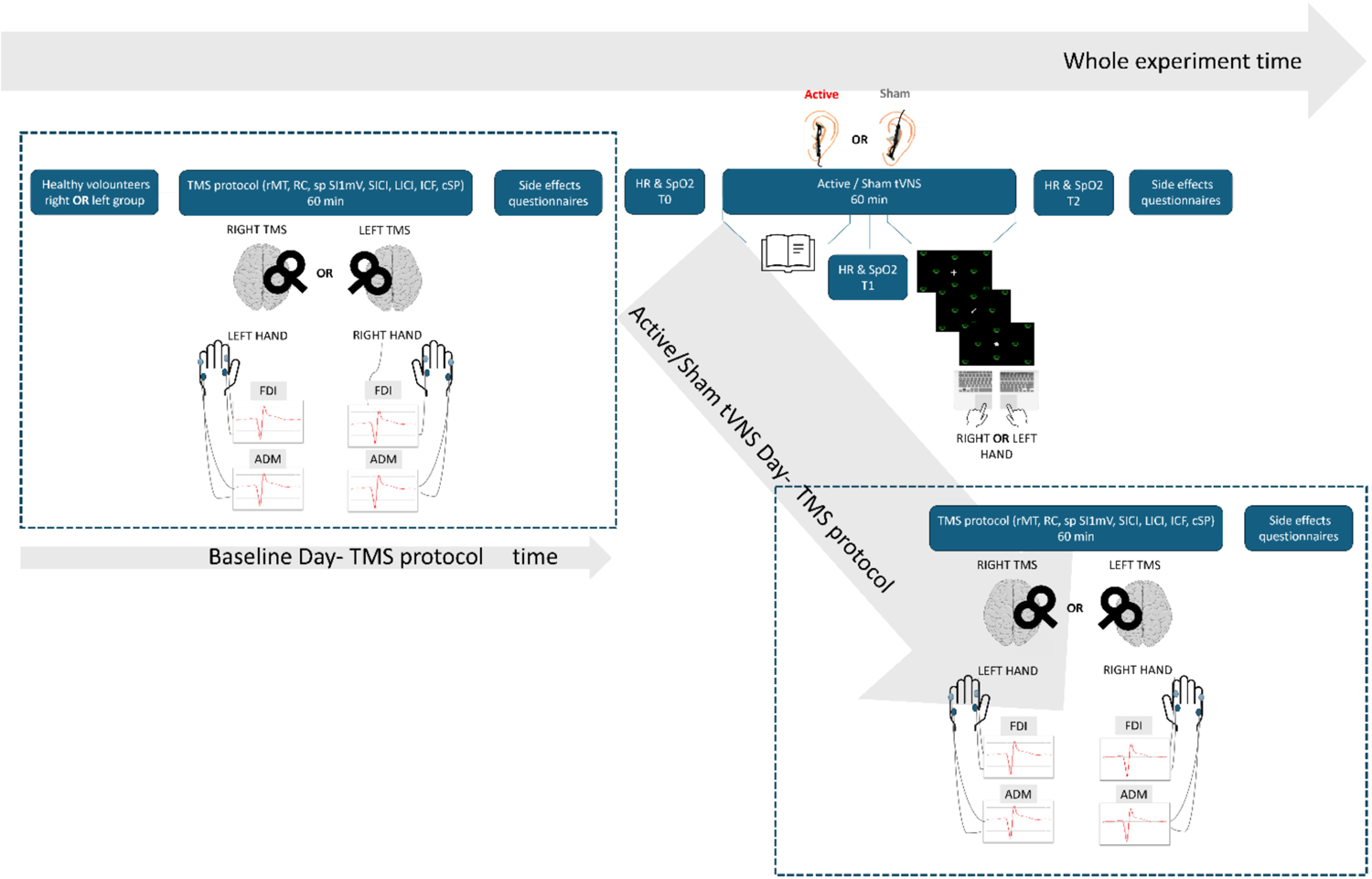

### TMS protocol

Single pulse and paired pulse TMS were delivered to scalp portion overlying the left (left TMS group) or the right (right TMS) motor hand region through a 70-mm-figure of eight coil (Magstim polyurethane-coated coil) connected to a Magstim 200^2^ BiStim stimulator (Magstim Company, Carmarthenshire, Wales, UK).

Participants were seated on a comfortable recliner chair with their arms resting on a pillow. They were asked to keep their hands as still and relaxed as possible. After cleaning with alcohol the skin of the hand contralateral the stimulated motor cortex, we applied 1 cm Silver/Silver Chloride cup electrodes (SEI EMG srl) to record the electromyographic (EMG) activation from the First Dorsal Interossei (FDI) and the Abductor Digiti Minimi (ADM) muscles. The electrodes were placed in a belly-tendon configuration. Electrodes position for the FDI and the ADM muscles was determined, in two separate moments for each muscle, by palpation during maximal voluntary contraction (i.e., the abduction of the index finger or the abduction of the little finger while the experimenter exerted a pressure toward the opposite direction). For each muscle a cathodic electrode was placed over the corresponding metacarpal phalangeal joint. Finally, the ground electrode was located over the ventral part of the wrist. Next, all the electrodes were connected to a Biopac M-160 system (BIOPAC Systems, Inc., Goleta, CA) allowing the amplification (gain: 1000 mv), online band pass filtering (5Hz to 2 kHz, notch filter at 50 Hz) and digitization of the EMG signal (sampling rate: 6250 kHz). The signal was stored on a personal computer for data display and later offline data analyses. The EMG data were collected for 300 ms starting 100 ms before the TMS pulse. After the electrode montage, participants were asked to wear a swim cap, allowing us to mark the coil position after the determination of the motor hotspot and to ensure the same coil placement throughout the three sessions. The motor hotspot for the activation of both muscles (i.e., the scalp position from which maximal-amplitude MEPs were elicited) was determined by moving the coil in approximately 0.5 cm steps around the presumed motor area and stimulating with a constant, suprathreshold stimulus intensity. The coil was held tangentially to the scalp with the handle pointing backward and laterally to form a 45° angle with the sagittal plane. This coil orientation induced a posterior-anterior current in the brain. This position was then marked on the cap and it was kept constant during the following MEP recording. Then, the recording session started with the following measures. First, the resting Motor Threshold (rMT) (which was expressed as percentage of maximum stimulator output) was determined; rMT was defined as the minimum stimulus intensity (SI) able to produce MEPs of at least 0.05 mV peak-to-peak amplitude in at least the 50% of the trials at rest in the lowest threshold muscle (the FDI). This procedure was used to avoid saturation of Corticospinal Excitability (CSE) modulation (Devanne et al., 1997). When a plausible value of rMT was approached, this value was confirmed when at least 8 MEPs were obtained after delivering spTMS for 16 trials every ∼8 seconds.

We then determined the Recruitment Curve of EMG responses by recording MEPs at five increasing TMS intensities. MEPs were elicited at SI corresponding to the 110%, 120%, 130%, 140%, and 150% of the rMT. Ten pulses were delivered for each SI every 8 seconds. On the basis of the peak-to-peak amplitude of the MEPs recorded during the RC, we selected for each participant the SI at which we were able to evoke MEPs at rest of approximately 1 mv in the FDI muscle (SI_1mV_). This intensity was used as suprathreshold SI while the intensity corresponding to the 80% of the rMT was used as subthreshold conditioning SI (SI_80%_). In particular, in the single pulse SI_1mV_ recording, single pulse TMS was delivered at the SI_1mV_ intensity. For SICI recording, the conditioning SI_80%_ TMS pulse and the test SI_1mV_ stimulus were delivered with an interstimulus interval (ISI) of 3 ms. For LICI recording, the test and the conditioning SI_1mV_ pulses were delivered at an ISI of 100 ms. For the ICF, a conditioning SI_80%_ pulse was followed with an ISI of 10 ms by a test SI_1mV_ pulse. In all cases, the test stimuli were separated by at least 10 seconds, and we varied the duration of the fixation cross before and after the TMS pulse to reduce expectations about the TMS delivery. For each recording, 16 pulses or pairs of pulses (for pp-protocols) were delivered. The order between the four recordings (single pulse SI_1mV_, SICI, LICI, ICF) was counterbalanced between participants but it was kept constant across sessions.

Lastly, the cortical silent period (cSP) was recorded. Participants were firstly instructed to perform a maximal active contraction of the FDI muscle by opposing their index finger against the thumb of the experimenter and by maintaining this maximum voluntary contraction for 30 seconds. Then, in the following 30 seconds they were asked to reduce the force to the 30% of their maximum capability. Continuous EMG activity was recorded during this process. Immediately afterward, 16 pulses at 120% of the rMT were delivered every ∼8 seconds while participants continued to contract their index finger at 30% of their maximum strength (see Fig. 2). In this case, EMG was collected for 1,300 ms starting 300 ms before the TMS pulse. The timing of EMG recording and TMS triggering was controlled with the E-Prime 3.0 software (Psychology Software Tools Inc., Pittsburg, PA) running on a pc.

**Figure 2:**
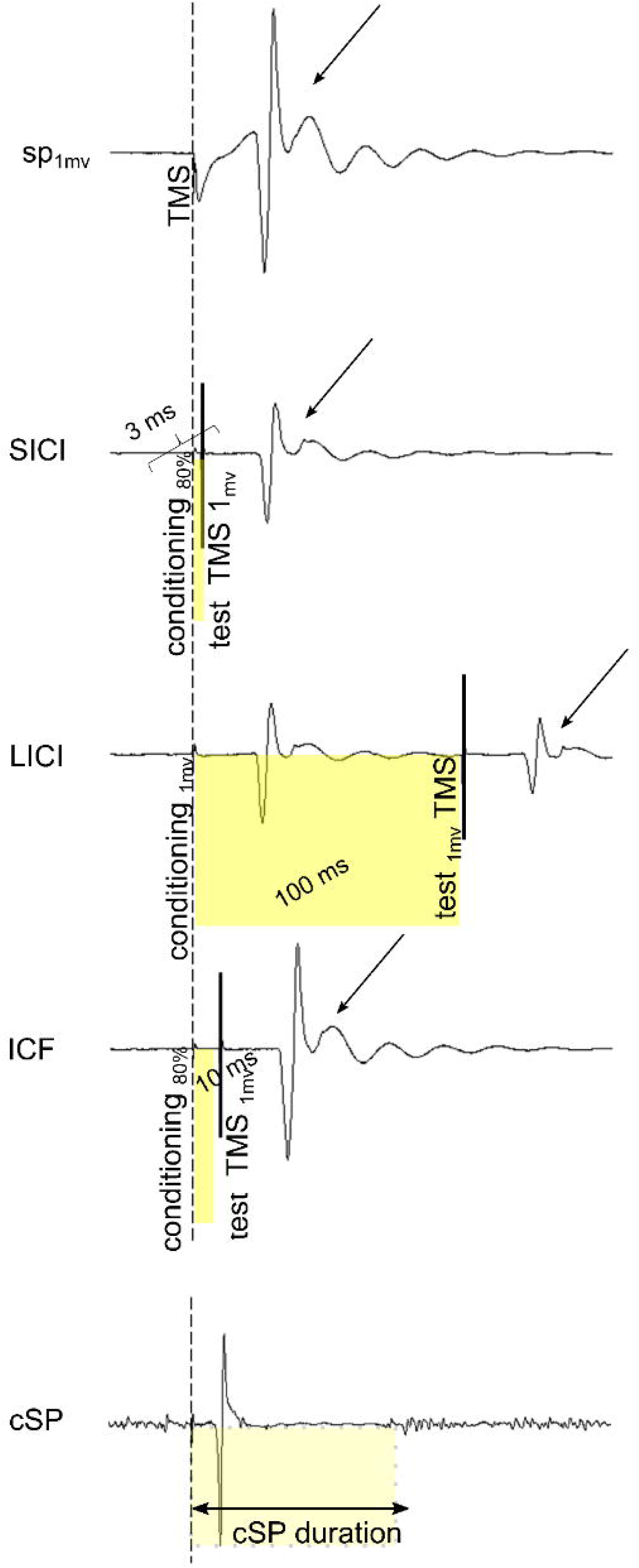

The TMS protocol, lasting approximately 60 minutes, was repeated in the following two sessions soon after the end of tVNS. The structure of the TMS protocol was kept identical across sessions and it was counterbalanced between participants.

### tVNS

TVNS was performed using the tVNS-E ® device (tVNS Technologies GmbH), composed by a programmable stimulation unit connect to two titan ear electrodes that are mounted on a gel frame. This allows generating and transferring electric pulses from the stimulation unit to the surface of the skin, where the electrodes are applied. After asking participants to clean their left external ear with alcohol swabs, we used a spray solution over the electrodes to enhance conductivity. For active tVNS condition, the electrodes were applied over the Cymba conchae, i.e., the region of the ear that has been shown to contain the highest density of ABVN projections (Peuker & Filler, 2002). The sham stimulation was provided by placing the electrodes over the earlobe, an ear target with minimal innervation from the vagus (Peuker & Filler, 2002). In this way, both tVNS stimulation conditions were able to induce reliable sensations but only the active tVNS electrode placement configuration activated the vagus nerve pathway. Given that some of the efferent fibers in the right vagus nerve regulate heart rate (Nemeroff et al., 2006), electrodes were placed on the left ear only, whose stimulation does not exert arrhythmic effects (Kreuzer et al., 2012). The intensity of tVNS was set at a level corresponding to the perceptual threshold, i.e., at the intensity in which participants could perceived sensations and 0.1 mA below the intensity at which they reported discomfort at the electrode site. This method was used for both the active and the sham sessions. Stimulation alternated between On/Off periods of 30s each, with a frequency of 25Hz, pulse width of 250 µsec. tVNS was applied for a total of 60 minutes, in each session.

### Visuomotor task

Contingent with the administration of tVNS, an ad hoc computer-based task was administered. The task was built to evaluate manual visuomotor control and it was programmed in E-Prime 3 software (Psychology Software Tools, Inc., Pittsburgh, PA, USA) and presented on a 17-inch monitor with a refresh rate of 60 Hz and a resolution of 1280×768.

During this task, participants performed a click-and-drag operation on a crumpled-paper object appearing at the center of the screen. The goal was to drag and drop this object to a location indicated by a previously displayed arrow pointing towards a target object (a recycling bin) within a configuration of objects. Participants focused on the arrow’s direction to accurately move the crumpled paper to the designated target bin, using the touchpad by using their right or left index finger. They were instructed to perform the task only by using the index finger. The use of the left or the right hand to perform the task was determined by the participants’ allocation to the TMS-site group: those stimulated over the right M1 used their left hand, and those stimulated over the left M1 used their right one.

Each experimental trial began with a fixation cross displayed for 2000ms, followed by an arrow appearing at the center of the screen for 500ms. Subsequently, the object appeared at the center of the screen. Participants were instructed to complete the click-and-drag operation within 2500ms. Participants were allowed to click and collect the object within 1000ms and to complete the click-and-drag operation within 1500 ms; then the object disappeared (Fig. 3).

**Figure 3:**
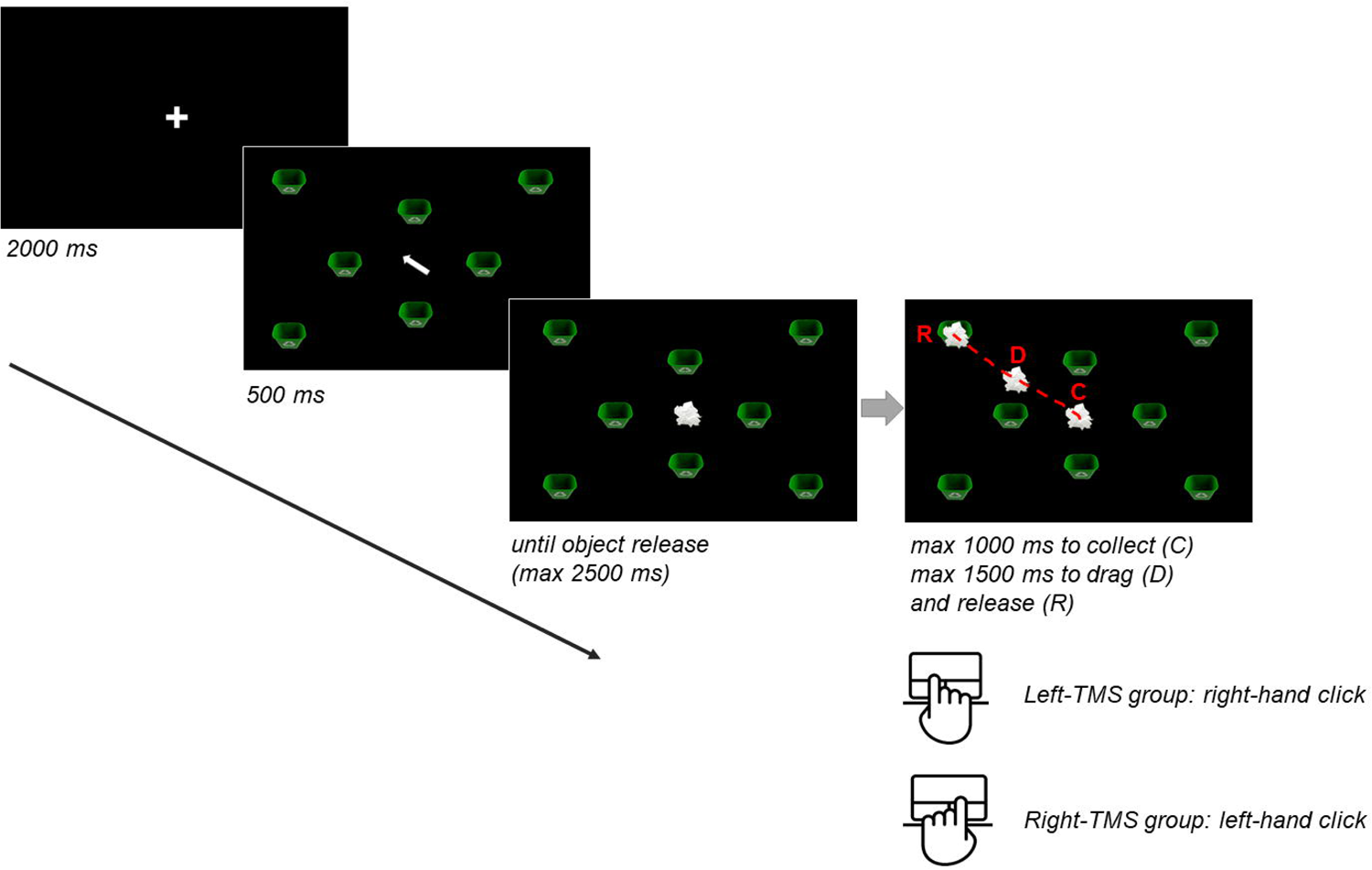

Two distinct configurations were created, each displaying eight recycling bins: four positioned near the center of the screen (“near” distance) and four positioned further away (“far” distance) (Fig. 4A). The locations of the recycling bins were determined using percentage values, as default setting (Fig. 4B).

**Figure 4:**
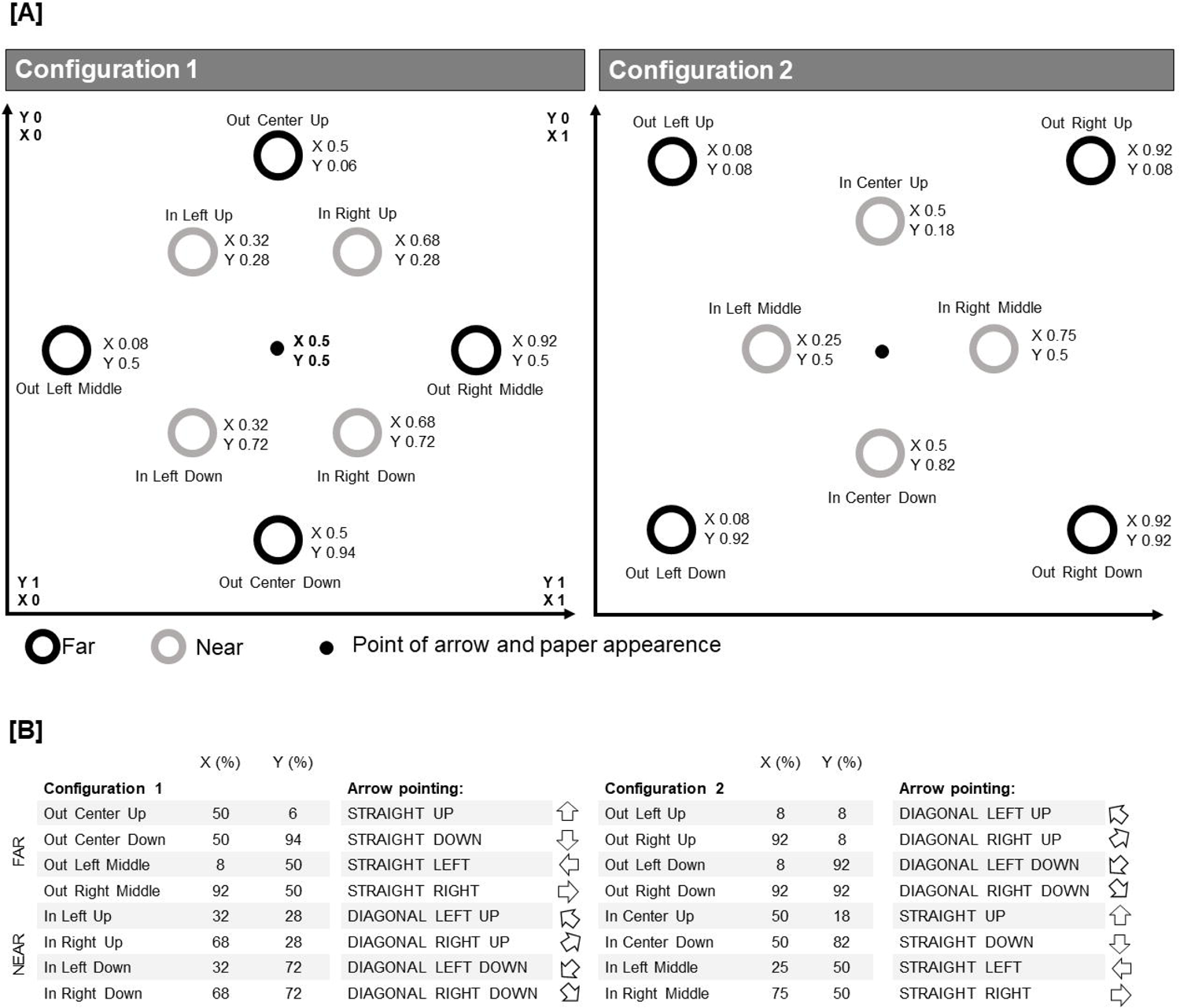

Participants completed four blocks of 80 trials each, totaling 320 trials. Each configuration accounted for 50% of the trials, with their presentation randomized within blocks (i.e., 40 trials per configuration).

### Safety and tolerability evaluation

With respect to safety, to check for unwanted effects of tVNS, heart Rate (HR) and oxygen saturation (SpO2) were recorded, by using a wrist pulse oximeter (GIMA 34340, GIMA Spa, Italy). The device was applied on the left hand for all participants. Cardiovascular parameters were recorded during a two-minute rest period at three time points: before the start of tVNS (T0), 30 minutes after the start of stimulation (T1), and at the end of the tVNS session (T2). For each measurement, 15 values were recorded, every 8 seconds.

With respect to tolerability, after the end of each tVNS stimulation day, participants were asked to fill-in a questionnaire assessing the sensation and side effects caused by tVNS, by adapting the questionnaire by Fertonani and colleagues (Fertonani et al., 2015). In particular we asked participants to report the intensity of each possible perceptual sensation (burning, fatigue, heat, pain, itching, iron taste, tingling) using a 5-point Likert scale, and to report any other eventually experienced sensations. At the end of each session, they were also asked to report the sensations experienced during the application of the TMS protocols by using an adapted version of the questionnaire by Giustiniani and colleagues (Giustiniani et al., 2022).

### Data handling and statistical analysis

Statistical analyses were performed using STATISTICA 8.0 (StatSoftInc, Tulsa, Oklahoma) and R software. Data are reported as mean (M) ± standard error of the mean (SEM). The level of statistical significance was set to p < 0.05 and effect sizes were estimated using partial eta squared (η_p_^2^). To follow-up significant interactions, Duncan post hoc tests were performed to reduce the risk of false negative (Type II) error when correcting for multiple comparison. The significance threshold was set at 0.05 for all statistical tests.

### Visuomotor task

Trials in which the participant successfully completed the click-and-drag operation within the specified time constraints were labeled as “valid”. If the participant failed to click and collect the paper within 1000ms, the trial was labeled as “invalid” and subsequently discarded from the analysis. If the participant correctly initiated the drag operation, but failed to release the paper within 1500ms, the program recorded the final position of the paper before starting a new trial. These trials were labeled as “overtime”. The task measured three metrics: precision error (PE), and reaction time (RT). PE was calculated as the Euclidean distance between the drop position of the paper and the actual cued bin position. To compute PE, the x and y coordinates of the bins (bin_x_, bin_y_) and the final drop position of the paper (drop_x_, drop_y_) were converted to pixels based on the screen dimension (e.g., if the x coordinate of a final drop position was recorded as 7% of the screen, its position in pixel was (7 / 100) * 1280 = 89.6). The new pixel values were then translated into a Cartesian plane with the origin at the top-left corner (Figure 4A; see Configuration 1), dividing the newly transformed x coordinates by 1280 and the y coordinates by 768. The Euclidean distance between the target bin and the final drop position of the paper in each trial was then calculated using the formula: 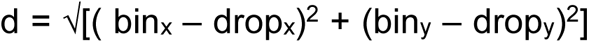. The lower the distance between the target bin and the drop position of the paper (values approaching 0), the more precise the release operation. RT, measured in milliseconds, was defined as the interval from the appearance of the paper to the click-to-collect operation. This metric was used to assess readiness to respond.

Invalid trials (7.9% of the total data) in which participants failed in collecting the data within 1000ms were discarded from analysis. Next, three separated mixed ANOVA were conducted on PE and RT. All models featured a 2 × 2 × 4 × 2 × 2 structure, including the following factors: Stimulation (active vs. sham), Distance (distance of the target bin from the central object: near vs. far), and Block (1, 2, 3, 4) as within-subject factors, and Hand (task performed with right vs. left hand) and Gender (female vs. male) as between-subject factors. The analysis on PE included both valid and overtime trials, whereas the analysis on RT included only valid trials, where participants successfully performed the drag-to-drop operation within the time limits. After the removal of invalid trials, data was filtered based on RT using a robust method for outlier detection (Lachaud & Renaud, 2011). We used the median as a robust estimate of the center (rob-center) and the tau-estimate as a robust measure of scale (rob-scale) (Maronna & Zamar, 2002). Unlike the arithmetic mean and standard deviation, which are heavily influenced by outliers, this method provides more reliable estimates in their presence. Trials with RT falling below −2.5 rob-scale units from the rob-center value, calculated within each participant, block, stimulation and distance conditions, were removed. The procedure led to the removal of 98 trials (0.69%) in the active stimulation condition and 103 trials (0.73%) in the sham stimulation condition. Overtime trials made up 3.4% of the data included in the analysis, and valid trials accounted for 96.6%.

### TMS indices

With respect to MEP processing, we extracted peak-to-peak amplitude (expressed in mv) of MEP recorded in each condition for the two muscles. To ensure that MEPs were acquired under full muscle relaxation (Devanne et al., 1997), MEPs were discarded if the EMG signal showed pre-TMS EMG activity. Thus, background pre-TMS EMG activity was computed for each trial as peak-to-peak amplitude in the 70-ms window before the TMS pulse. MEP with an EMG background deviating more than 2 SD from the mean were removed from further analysis. Further, we excluded trials with MEP amplitudes lower than 0.05 mv. MEP amplitudes values were then averaged for each measurement, separately for each participant and for the two muscles, and used for computing the indices described below.

Cortico-spinal excitability. Cortico-spinal excitability was indexed by rMT and recruitment curves. With respect to the recruitment curves, MEP amplitudes were averaged separately for each participant, muscle, day of stimulation and intensity of stimulation. Thus, for each participant, muscle and day of stimulation, the unstandardized beta coefficient (herein after beta index) was calculated across trials using regression analysis with intensity of stimulation (110%, 120%, 130%, 140%, 150%) and MEP amplitude as the independent and dependent variables, respectively. This index would represent the steepness of the regression line between the 5 levels of Intensity of Stimulation, representing the Input-Output curve slope.

Intracortical inhibition. SICI and LICI values were expressed as the ratio between the mean conditioned MEPs amplitude in the SICI and LICI recordings, respectively, and the mean single pulse SI_1mV_ MEPs amplitude (mean MEPs condition/mean Single pulse SI_1mV_ MEPs), with smaller values suggesting greater inhibition. Silent period durations (cSP) was measured for each trial as the time between the delivery of the TMS pulse and the return of EMG activity by visual inspection.

Intracortical facilitation (ICF). As for SICI and LICI, ICF was computed as the ratio between the mean conditioned MEPs amplitude in the ICF recording and the mean single pulse mean single pulse SI_1mV_ MEPs amplitude (mean MEPs condition/ mean Single pulse SI_1mV_ MEPs), with higher values suggesting stronger facilitation.

All the indexes, except cSP, were submitted to mixed ANOVAs, with Group and Gender as between-subject factors and muscle and day of stimulation as within-subject variables. The analysis of cSP included Group and Gender as between subject factors and the day of stimulation as within subject variable, since, for this index, only the FDI muscle was considered, being the muscle involved in active contraction. Due to technical problems during data acquisition, the analyses of recruitment curves and cSP were conducted on 48 participants (28 female). Furthermore, due to the known variability in individuals’ responses to pp-TMS paradigms several participants did not show any visible MEPs in response to the conditioned TMS pulse during the recording of intracortical inhibition and/or facilitation indexes, particularly at longer ISI; since SICI, LICI or ICF could not be calculated, their data were excluded from the relative analysis. Consequently, the analyses were conducted on a sample of 43 participants (25 female) for the SICI index, on 24 participants for LICI (16 female), and on 39 participants (20 female) for ICF.

### Safety measures

HR and SPO2 data were entered into two separated ANOVAs. We run two separated 2×3×2×2 rmANOVA with tVNS stimulation (active and sham) and time of recordings (t0: beginning of experiment, t1: after 30 minutes of tVNS, t2: after 60 minutes of tVNS, after at the end of tVNS) as within-subject variables, while gender (male or female) and TMS stimulation group (right or left) were entered in the model as between-subject variables.

### tVNS side effects

To explore potential differences in ratings of tVNS-induced sensations between active and sham stimulation conditions, a series of comparisons were conducted for each sensation by means of the Wilcoxon Matched Paired Test.

## Results

### Behavioural results of the visuomotor task

PE. The ANOVA showed a main effect of Distance (F_1,44_ = 114.6, p <.00001; η_p_^2^ = 0.72), as participants performed more precise drop-operation (i.e., decreased PE) when the target bin was near (0.12 ± 0.001) as compared to when it was far from the central object (0.14 ± 0.001). The model also yielded a significant interaction of Stimulation * Block * Hand (F_3,132_ = 3.07, p =.03, η_p_^2^ = 0.07). Post-hoc tests showed that, only in participants who performed the task with the right hand, the precision of the drop-operation significantly improved in the fourth block (0.12 ± 0.002) as compared to both the third (0.13 ± 0.002; p =.01) and the second block (0.13 ± 0.003; p =.03) during active stimulation (see Fig. 5). The analysis did not yield any other significant effects (all F < 3, all ps >.07).

**Figure 5:**
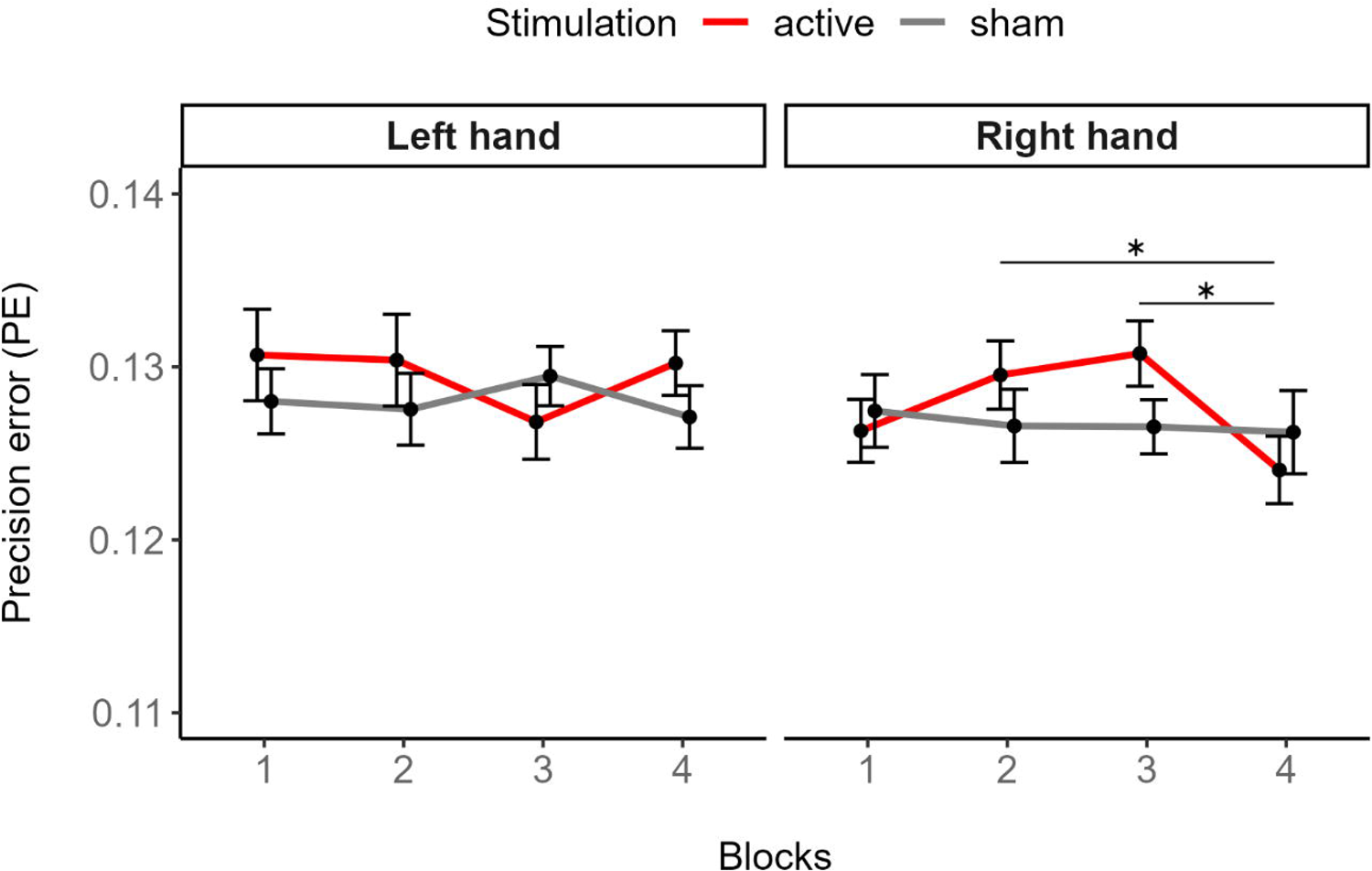

RT. The analysis on RT showed a significant main effect of Block (F_3,132_ = 10.07, p <.00001; η_p_^2^ = 0.19), as the click-to-collect reaction times decreased from the first block (382.75 ± 10.95) to the second (368.20 ± 12.38; p = .02) and third block (351.81 ± 11.64; p <.00001). Moreover, RT was significantly faster in the third than in the second block (p = .01). Likewise, the click-to-collect RT in the fourth block (350.72 ± 12.89) was faster than that recorded in the first (p <.00001) and second block (p = .01), whereas the difference between the third and fourth block was not statistically significant (p = .86). The model did not show any other significant effect (all F < 3, all ps >.05).

### TMS index

#### Cortico-spinal excitability

rMT. The analyses of the rMT revealed a significant main effect of Stimulation day (F_2,90_=14.501; p<.001; 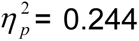), with lower rMT during the baseline (34.673%± 0.578) compared to either the Active (35.980% ± 0.593; p<.001) or the Sham (35.714% ± 0.634, p<.001; see Figure 5) stimulation day, without differences between the two tVNS days (p=.273). No other significant main effect or interactions were observed (all ps> .392, Fig. 6).

**Figure 6:**
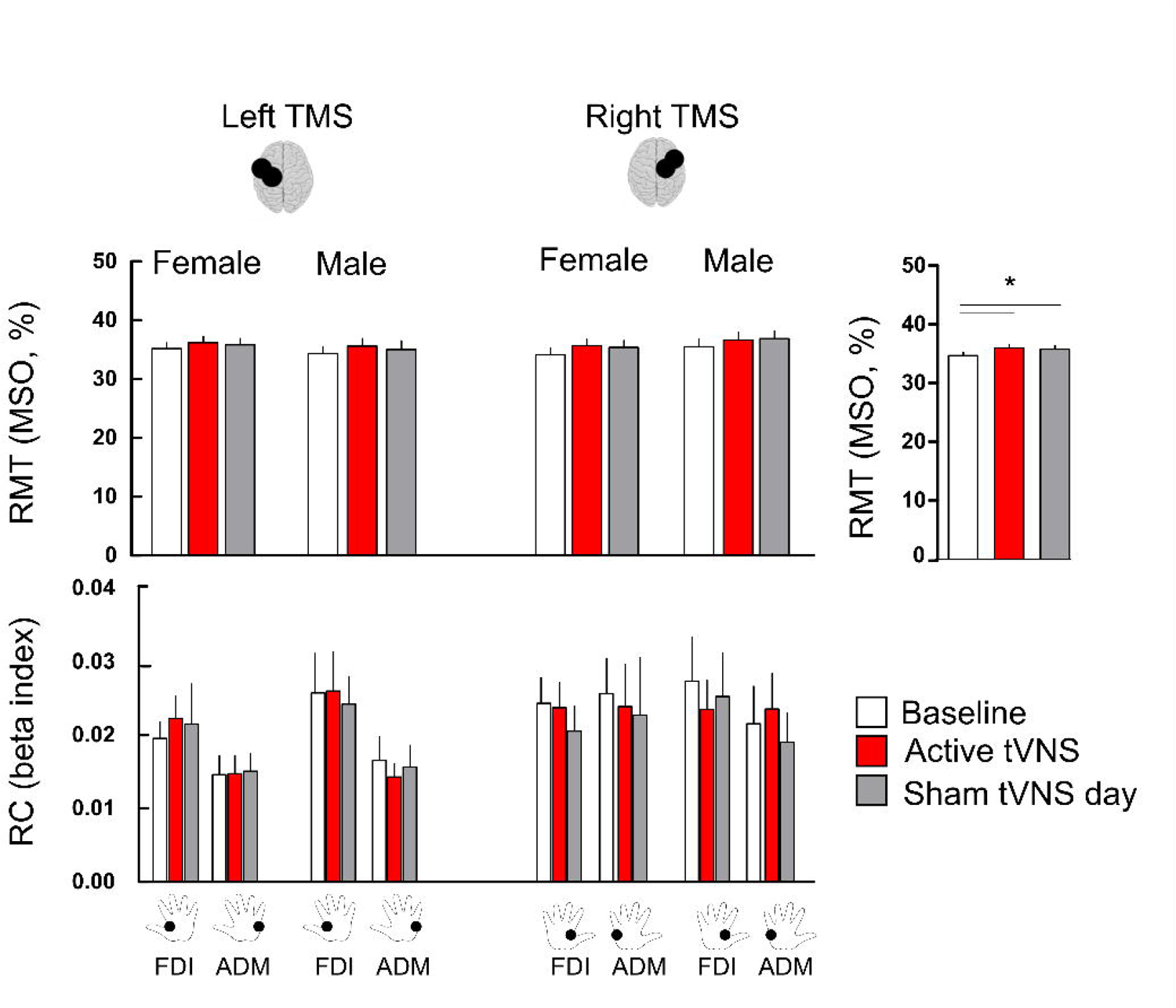

Recruitment curves. A tendency toward significance emerged for the main effect of Muscle (F_1,44_=3.961; p=.053; 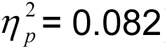), due to a steeper increase of the curve for the FDI than the ADM muscle (FDI: 0.023± 0.001; ADM: 0.019 ± 0.001). No other significant main effects or interactions were found (all ps > .178, Fig. 6).

#### Cortical inhibition

SICI. No main effects were significant (all p>.381). However, we found a significant three-way Muscle × Stimulation day × Gender interaction (F_2,78_ = 6.798; p=.002; 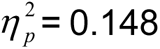) which was further qualified by the TMS Group factor in the four-way Muscle × Stimulation day × Gender × TMS Group Interaction (F_2,78_ = 3.477; p=.035; 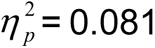, Fig. 7). To explore this interaction, two different mixed-model ANOVAs were performed, for the male and the female gender group with the TMS Group as between-subject factor and Muscle and Day of stimulation as within-subject variables. With respect to the male group, no main effects nor interactions were significant (all ps> .085). Differently, in the female group, we observed a significant Muscle × Stimulation day interaction (F_2,46_ =5.167; p= .009; 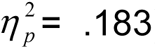). Interestingly, post hoc comparison revealed that, for the FDI muscle, the index was smaller in the Active (0.209± 0.018) with respect to either the Baseline (0.285 0.030, p=.009) or the Sham stimulation day (0.272 ± 0.030; p= .019), which did not differ between each other (p= .645); this indicates a selective increase of GABA-A mediated cortical inhibition after active tVNS. Of note, the difference with respect to the ADM muscle in the Active stimulation day was also significant, due to smaller values for the FDI than for the ADM (Active, ADM= .311 ± 0.046, p <.001). No other main effect or interactions emerged as significant (all ps> .124).

**Figure 7:**
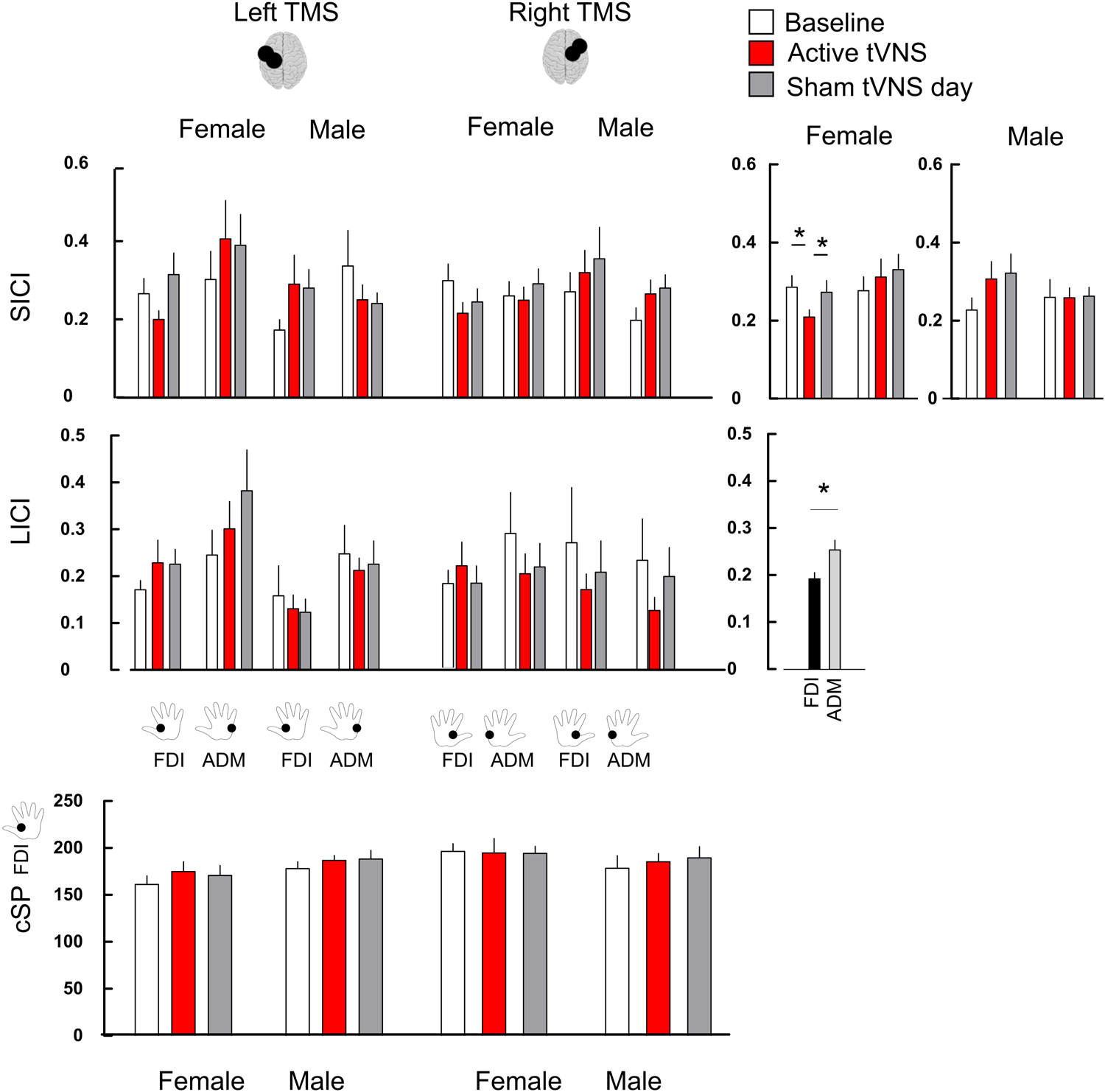

LICI. The main effect of muscle was significant (F_1,21_ =4.893; p= .038; 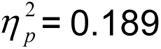) due to smaller values of this index (suggesting stronger inhibition) for the FDI (0.192 ± 0.133) than the ADM (0.253 ± 0.017). No other significant main effect or interaction emerged (all ps> 0.066, Fig.7).

cSP. No main effect or significant interaction were found on the cSP duration (all ps> .119; Fig. 7).

#### Intracortical facilitation

ICF. The Muscle × Gender × Group Interaction was significant (F_1,35_=6.610; p=.015; 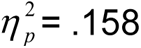). However, post-hoc comparisons did not show any significance (all ps> .291). No other significant main effect or interaction emerged (all ps> .228, Fig. 8).

**Figure 8:**
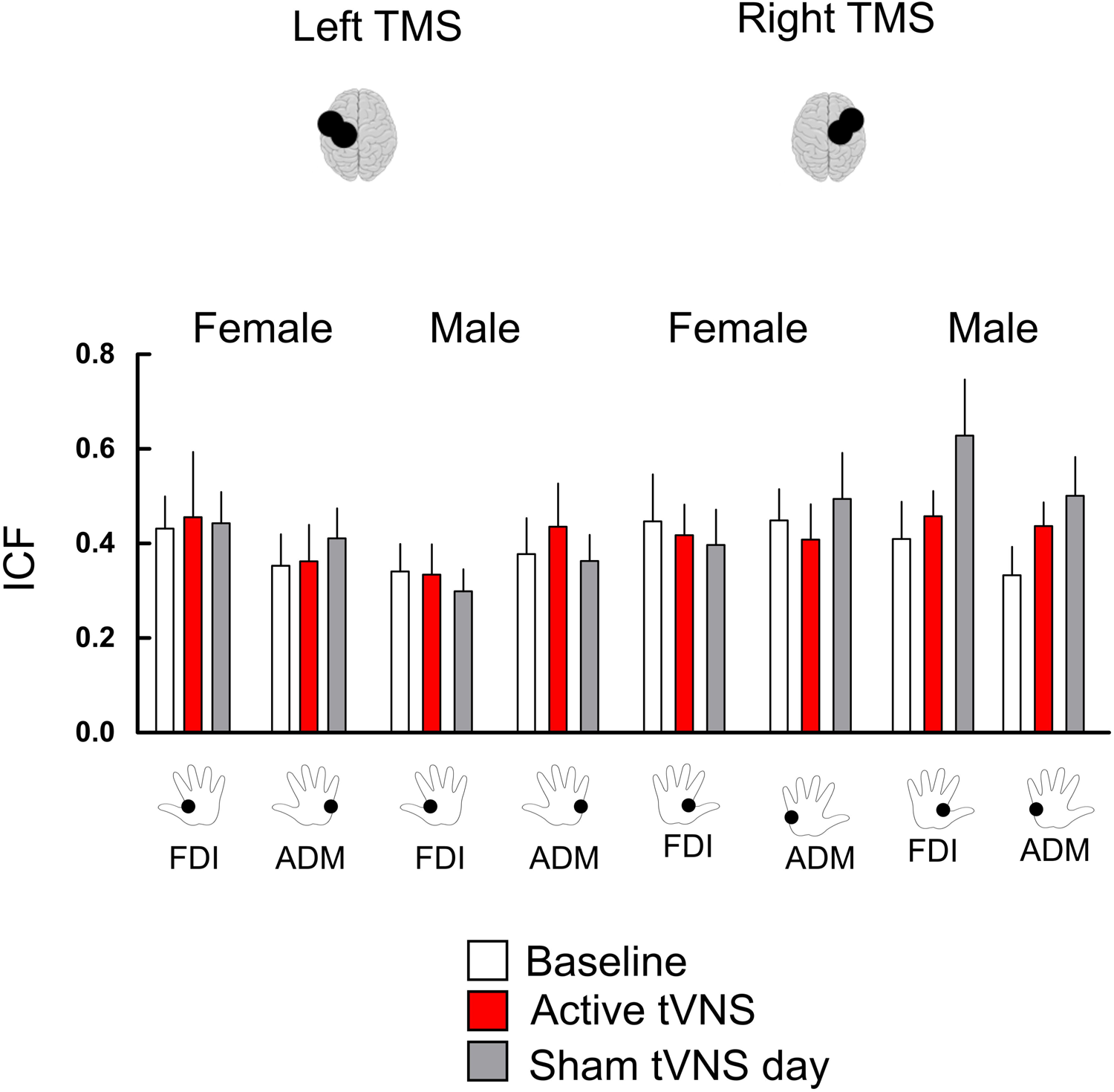

#### Safety measures

HR. Analysis revealed a main effect of Gender (F_₁,₄₅_= 4.232, p= 0.045, 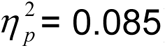), as females had higher HR (73.27 ± 1.97) compared to males (66.81 ± 2.53). Also, a significant main effect of time emerged (F_₂, ₉₀_= 33.472, p< 0.001, 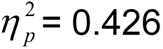), described by a progressive decrease of HR over each recording time (t0= 74.05 ± 2.72; t1= 69.87 ± 2.10; t2= 67.98 ± 1.93). No other significant effects emerged (all ps> 0.066).

SPO2. Analysis on SPO2 highlighted a main effect of Gender (F_₁,₄₄_= 4.4, p= 0.0422, 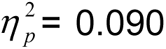), as females displayed higher SPO2 level (97.88 ± 0.37) than males (96.22 ± 0.43). No other significant effects emerged (all ps> 0.154).

#### tVNS side effects

Analysis revealed that both pain (z_12_ = 3.059, p =.002) and burning sensation (z_11_ = 2.044, p =.040) sensations were higher during Active (pain: M = .411 ± .10; burning: M = .313 ± .09) compared to Sham stimulation (pain: M = .078 ± .047; burning: M = .137 ± .056). However, participants did not report a significant difference in how much they thought these sensations affected their performance in the visuomotor task between the two conditions (z = 0.258, p =.795) (active: M = .275 ± .075; sham: M = .314 ± .091), suggesting that an effective blinding was achieved. No other significant differences were found (all ps >.123). Finally, all participants referred that they perceived as equally effective both stimulation conditions, suggesting that they were blinded to the manipulation until the end of the whole experiment.

## Discussion

The stimulation of the Vagus Nerve has been proposed as a bottom-up track to modulate neural activity and promote neuroplasticity. Indeed, the activation of the Vagus Nerve is thought to propagate to subcortical and cortical brain structures, enhancing brain GABA (Marrosu et al., 2003) and Noradrenaline levels, which play a pivotal role in brain functioning and plasticity. While most of the studies highlighted the effect on the noradrenergic system (Berger et al., 2021), the effects on GABA transmission are less clear. With this respect, animal and clinical studies with invasive VNS demonstrated that long term stimulation increases free GABA in cerebrospinal fluid of patients with epilepsy, with more favourably effects of VNS in patients with higher GABAa receptor density (Ben-Menachem et al., 1995; Hammond et al., 1992; Marrosu et al., 2003). Despite behavioral evidence pointing to an effect of tVNS on motor inhibition, the neurophysiological effects and mechanisms of tVNS on GABA-ergic activity are less clear.

By combining sp- and pp-TMS of left and right motor cortex in male and female participants, here we tested the effects of tVNS on neurophysiological measures of CSE, intracortical inhibition, and ICF as well as on visuomotor performance. We found that tVNS specifically increased intracortical inhibition mediated by GABAa activity, while leaving unaffected CSE, ICF and intracortical inhibition mediated by GABAb activity. It also improved visuomotor learning. These effects, however, were moderated by the site of the stimulated motor cortex and by the gender of the participants. Indeed, left-ear tVNS increased SICI in both hemispheres, thus for MEPs recorded from both the left and the right hands, but only in female participants. Furthermore, the improvement in visuomotor learning performance only occurred in the participants, of both gender, that executed the task with the dominant right hand (i.e., in the left TMS group). The results add to previous studies (Capone et al., 2015; Midden et al., 2023; Mertens et al., 2022; Gerges et al., 2024; Yun et al., 2024) by providing supportive evidence on the effects of tVNS on GABA activity, but specifying the effects in terms of participants’ gender and laterality.

The laterality of the effects of left-ear tVNS have been poorly investigated in previous studies. In keeping with the effects documented after invasive VNS (Di Lazzaro et al., 2004), Capone et al. (2005) examined the changes in CSE and SICI in the right and the left motor cortex after 60-min of left tVNS. They found that tVNS increased SICI, but not CSE, in the contralateral (right) motor cortex, suggesting an increase in GABA (and in particular GABAa) concentration (Stagg et al., 2011). Unfortunately, the sample size of that study was limited to 10 participants (7 females). Moreover, tVNS was administered on the tragus, whose stimulation is doubtful inducing optimal and specific vagal activation (Peuker & Filler, 2002). More recently, van Midden et al. (Midden et al., 2023) replicated the study with a larger sample size of 30 participants (17 females) by targeting the cymba concha for tVNS. Importantly, the cymba conchae seems to produce stronger activation of the vagal nerve with respect to tragus (Yakunina et al., 2017). Their findings showed an increase in right-motor cortex SICI, but no change of CSE, after active-compared to sham-tVNS.

Notably, both Capone (Capone et al., 2015) and van Midden (Midden et al., 2023) studies focused solely on GABAa receptor-mediated inhibition. To explore a possible effect on GABAb receptor-mediated inhibition, Mertens et al. (2022) and Greges et al., (2024) evaluated the effects of 60 minutes of tVNS delivered in the left cymba concha on both GABAa and GABAb-mediated motor cortical inhibition indexes (i.e., SICI and LICI, respectively) as recorded from the contro-lateral right motor cortex. Mertens and colleagues (Mertens et al., 2022) found no tVNS-induced modulations on cortical GABAergic inhibition. However, in this last case, only 15 male participants were tested; if this allowed reducing the impact of hormonal confounding variables on cortical measures (Smith et al., 2002), it hampered the generalizability of the results (Yokota et al., 2022). Differently, the study by Greges and colleagues (2024) delved into the effects of tVNS in middle-aged and older male and female adults by comparing the effect of 60 minutes or 30 minutes of active stimulation with 30 minutes of sham stimulation. While no effects were observed on LICI, with respect to SICI the results corroborated the findings from Capone et al., (2015) and van Midden at al., (2023) of tVNS increasing inhibition. This effect was specifically limited to the 60 minutes of active stimulation, suggesting that varying doses of tVNS may yield different physiological outcomes. Providing further insight into the role of participant’s sex on the variability of tVNS effects, this effect was bigger in female participants, which represented the 65% of the whole tested sample. However, in this study the sham stimulation was delivered by switching off the stimulator. This did not allow ruling out that differences in sensations experienced during the active with respect to the sham condition could affect the results on cortical excitability and inhibition (Kojima et al., 2019; Luft et al., 2005; Sailer et al., 2002).

Based on the study by Capone and colleagues (2015) testing the effect of left tVNS on both left and right motor cortex and showing a selective effect on the right hemisphere, no other follow up studies investigated the issue of lateralization. In these cases (Gerges et al., 2024; Mertens et al., 2022; Midden et al., 2023), TMS protocols were indeed exclusively applied over the right motor cortex. The importance of a deeper investigation of lateralization was raised in the study by Keute and colleagues (2018). Indeed, by assuming a modulation of GABA transmission in the contralateral (right) motor cortex the authors primarily investigated the effects of tVNS on inhibitory processing to left hand responses. In particular, they investigated the effects of tVNS on a subliminal response priming paradigm that requires automatic motor inhibition. Their results indicated that left-ear tVNS modulated both the behavioral performance and the Event Related Potentials (ERP) components associated to inhibitory processes recorded in the contralateral (i.e., right) hemisphere. However, as the authors stated there was not enough evidence for the absence of a GABAergic effect in the ipsilateral hemisphere, and it was unclear whether the effects they reported (on the left hand) represented a rightward lateralization of the GABAergic effects of tVNS. By using a sham stimulation control, beside the baseline, no-stimulation condition, here we showed that left-ear tVNS increased SICI in both the ipsilateral and contralateral motor cortices, but this only occurred in female participants, thus providing explanatory evidence that may clarify previous discrepant findings.

The modulation of motor inhibition due to tVNS may be influenced by gender differences. Indeed, several studies have indicated that hormonal variations, particularly in relation to the menstrual cycle in females, could impact cortical inhibition (Smith et al., 1999, 2002; Zoghi et al., 2015). However, previous studies either exclusively recruited male participants (Mertens et al., 2022) or did not explore the impact of gender on their results (Capone et al., 2015; Van Midden et al., 2023; Yun et al., 2024). The fact that tVNS affects GABA activity in female participants must be considered in the use of tVNS as a treatment option. Beside differences in target area and participant’s gender, other issues as the uncontrolled state-dependent modulation of tVNS effects could account for disparities in results. The previous studies investigating the effects of tVNS on cortical inhibition (Capone et al., 2015; Gerges et al., 2024; Mertens et al., 2022; Midden et al., 2023) did not report what participants did during the delivery of tVNS, with the exception of Mertens et al., (2022) study, where participants could conduct relaxing activities during the intervention, while cognitive or physical intensive activities were not allowed. However, controlling for brain state, for instance by employing a task or by applying stimulation during a particular brain state, as done in this study, could contribute to reduce inter-individual differences in tVNS response.

By showing a state dependent effect of tVNS, the findings from this study could highlight also the importance of considering the timing of vagus stimulation with respect to the target examined and trained function (Capone et al., 2017; Khodaparast et al., 2013). Moreover, by pairing tVNS with the execution of a visuomotor task here we aim at resembling in a more ecological way the rehabilitative approaches combining standard motor rehabilitation with the application of tVNS, in order to foster and accelerate the outcomes of motor rehabilitative treatments (Capone et al., 2017).

We showed that tVNS not only increased GABAa-mediated cortical inhibition, but it also improved visuomotor performance, at least when the task was executed with the dominant right hand. GABAa receptor–mediated neurotransmission is particularly important to promote practice-dependent plasticity after brain injury and its alterations is considered an important patho-physiologic mechanism of motor dysfunction in many chronic conditions, such as stroke or cerebral palsy (Park et al., 2013). Accordingly, studies on animal models have demonstrated that invasive VN Stimulation (VNS) combined with motor practice promotes cortical reorganization, leading to greater improvement in forelimb motor recovery and increased synaptic motor connectivity as compared to motor practice alone (Meyers et al., 2018). In humans, a recent study (Dawson et al., 2021) has documented the beneficial effects (persisting after 90 days) of invasive VNS paired with physical rehabilitation, as compared to rehabilitation alone, in adults with moderate-to-severe arm impairment after chronic ischemic stroke. Unfortunately, invasive VNS necessitates a costly surgical procedure. The recent increase in clinical studies on tVNS for stroke patients underscores the attention this technique has garnered as a powerful tool for rehabilitating motor functional deficits (Badran et al., 2023; Baig et al., 2022; Capone et al., 2017), also paired with intensive bimanual training in children with Cerebral Palsy (NCT06372028, ClinicalTrials.gov Identifier).

The results of the current study support the possibility to exploit tVNS as a novel, non-invasive and bottom-up NIBS method to promote cortical plasticity after brain injury.

## Limitations

The conclusions that can be drawn from this study must be cautioned in front of some critical limitations. First, while we provided evidence for a muscle-specific modulation of tVNS for the muscle (i.e., FDI) that was involved in the visuomotor activity, the rMT and consequently the other parameters were set at the FDI muscle, being the lowest threshold one. Even if the procedure was used to avoid saturation of responses (Devanne et al., 1997), we could not exclude that this setting of parameters was suboptimal for the ADM muscle and thus that we were not able to catch the modulation also for this muscle. Furthermore, while we used a visuomotor task to exploit the possible state-dependency of tVNS effects, we did not have a condition in which participants were stimulated without performing any task or during performance of a different cognitive task. Thus, we cannot provide direct evidence on such state-dependency. Finally, generalizability to patient populations can be made only with caution. The present findings of state-dependent effects of tVNS may suggest that a patient’s brain with defective inhibitory mechanisms could be more or less prone to changes induced by tVNS as compared to healthy participants with normal cortical excitability measures.

