## Supplementary material for "Investigating the effects of transcutaneous Vagus Nerve Stimulation on motor cortex excitability and inhibition through paired-pulse Transcranial Magnetic Stimulation": Figure legends

**Figure 1.** Schematic representation of the experimental procedure. The first block (within the dotted line) represents the baseline day during which the TMS protocol was applied to measure cortical inhibition, facilitation and corticospinal excitability in a baseline condition at rest, when no tVNS was administered. On the right, the active and the sham tVNS stimulation days are represented. In these days, active or sham tVNS was administered during the execution of the visuomotor task, prior the TMS protocol. The procedure of the TMS protocol (represented within the dotted line) was repeated after tVNS as in the baseline stimulation day.

**Figure 2.** The arrows indicate the MEPs recorded in the sp1mv session, during SICI, LICI, ICF and the duration of the cSP. The vertical dotted line represents the single TMS pulse (for sp1mv and for cSP recording) and the conditioning TMS pulse (for SICI, LICI and ICF). The continuous black lines represent the test TMS pulse (for SICI, LICI and ICF). Different intervals between pulses are represented for the SICI (3 ms), LICI (100 ms) and ICF (10 ms), in yellow. The intensity of the first TMS pulse corresponded to the intensity able to evoke peak to peak 1mv MEP for the sp1mv session and the LICI session. For the SICI and the ICF it corresponded to the 80% of the rMT. For cSP the intensity of the TMS corresponded to the 120% of the rMT.

**Figure 3.** Example of experimental trial of the visuomotor task in which the participant had to drag-and-drop the paper in the bin located in the top-left corner of the screen, as signaled by the arrow. Depending on the TMS stimulation site, participants were required to right- or left-click using their index finger on the touchpad to collect (C) the paper within 1000 ms of its appearance. They then had to drag the paper towards (d) and release it above the bin (R) within 1500 ms from the collection operation.

**Figure 4.** Panel A: configurations of target objects (bins) and central objects (arrow and paper) along with their x and y coordinates translated into a Cartesian plane. Panel B: x and y coordinates of the bins, defined in percentages of the screen dimension, according to the configuration. Each bin, based on its location, was cued by a distinct arrow pointing towards it.

**Figure 5.** Mean PE across blocks (1,2,3,4) color-coded by stimulation condition (Active vs. Sham) for the Left (left panel) and right hand (right panel). Error bars represent ±1 SE. * $p < 0.05$.

**Figure 6.** Upper panel. Bar plot of rMT values separately for each group (female and male participants, left and right TMS group), in each day of stimulation (white: baseline day, red: active tVNS day, grey: sham tVNS day) as well as for the whole sample (on the right side) in the three stimulation days. Lower panel. Bar plot of the RC values (beta index) separately for each group (female and male participants, left and right TMS group), in each day of stimulation for the FDI and the ADM muscle. Error bars represent SE. * indicates significant effects at $p <.05$.

**Figure 7.** Upper panel. Bar plot of SICI values for the Left and the Right TMS group for the female and male Group, color-coded by Stimulation day (white: baseline day, red: active tVNS day, grey: sham tVNS day) for the FDI and ADM muscles (on the left side of the panel) and for the Female and Male group in the three stimulation days for the two muscles (in the right part of the panel). Central panel. Bar plot of LICI values for the Left and the Right TMS group for the female and male Group, color-coded by Stimulation for the two muscles (on the left side of panel) and by collapsing all conditions and groups, separately for the FDI (black) and ADM (grey) muscles (on the right side of the panel). Lower panel. Bar plot of CSP values of the FDI muscle for the Left and the Right TMS groups for the female and male Group, color-coded by Stimulation. Error bars represent SE. * $p < .05$.

**Figure 8.** Bar plot of ICF values for the Left and the Right TMS group for the female and male Group, color-coded by Stimulation (white: baseline day, red: active tVNS day, grey: sham tVNS day) for the two muscles.
